# Whole Tissue Imaging of Cellular Boundaries at Sub-Micron Resolutions for Automatic Cell Segmentation: Applications in Epithelial Bending of Ectodermal Appendages

**DOI:** 10.1101/2024.06.26.600880

**Authors:** Sam Norris, Jimmy K. Hu, Neil H Shubin

**Affiliations:** Department of Organismal Biology and Anatomy, The University of Chicago, Chicago, IL, USA; School of Dentistry, University of California Los Angeles, Los Angeles, California, USA; Molecular Biology Institute, University of California Los Angeles, Los Angeles, California, USA

## Abstract

For decades, biologists have relied on confocal microscopy to understand cellular morphology and fine details of tissue structure. However, traditional confocal microscopy of tissues faces limited light penetration, typically less than 100 µm, due to tissue opacity. To address this challenge, researchers have developed tissue clearing protocols compatible with confocal microscopy. Unfortunately, these protocols often struggle to retain cell boundary markers, especially at high resolutions necessary for precise cell segmentation. In this work, we introduce a method that preserves cell boundary markers and matches the refractive index of tissues with water. This technique enables the use of high-magnification, long working distance water-dipping objectives. The sub-micron resolutions achieved with this approach allows us to automatically segment each individual cell using a trained neural network segmentation model. These segmented images facilitate the quantification of cell properties and morphology of the entire three-dimensional tissue. As a demonstration of this methodology, we first examine mandibles of transgenic mice that express fluorescent proteins in their cell membranes. We then extend this technique to a non-model animal, the catshark, investigating the cellular properties of its dental lamina and dermal denticles – invaginating and evaginating ectodermal structures, respectively. Our technique thus provides a powerful tool to quantify in high throughput the 3D structures of cells and tissues during organ morphogenesis.

## Introduction

Biologists have long visualized the internal structure of cells and tissues by imaging thin sections with light or electron microscopy. These two-dimensional (2D) maps of biological information are analyzed to understand spatial relationships of RNA transcripts, proteins, cellular morphology, and tissue organization. Native tissue structures, however, are inherently three-dimensional (3D), justifying the need to move towards whole-tissue microscopy. Scientists have used traditional approaches such as MRI, x-ray computed tomography, and ultrasound imaging to acquire 3D information of tissues, but these techniques are unable to provide submicron resolutions, single molecule imaging, and lack certain contrasting methods compared to optical microscopy (*1*). Early methods to acquire 3D optical microscopy images include serial sectioning, but the resolution of the reconstructed 3D images is poor and limited by the slice thickness. Confocal and two-photon microscopy of fluorescently-tagged samples allow scientists to perform volumetric imaging at high resolutions (*2*), however, native tissues pose a complicated problem for optical microscopes: their composition of various lipids, proteins, minerals, pigments, among others components, render the tissues optically opaque due to refractive index mismatches, thus, scattering and distorting light, which limits the optical imaging depth of tissues (*3*).

Efforts to increase the transparency of tissues have been ongoing for at least a century, where early attempts focused on matching the refractive index (RI) of the surrounding medium to the RI of the tissue, which was further improved by bleaching, decalcifying, dehydrating, and removing hydrophilic components from the tissues (*4*). In the past decade, tissue clearing methods have accelerated: researchers have utilized a combination of advanced chemical techniques to remove high RI particles, such as lipids and fibrous proteins, endogenous pigments with strong absorption, such as heme and melanin, and the use of more efficient and appropriate RI-matching chemical agents without damaging or altering the native cell and tissue structures of interest (*3*). The combination of tissue clearing, fluorescence labeling (small-molecule dyes, tagged antibodies and fluorescent proteins), and efficient confocal, two-photon, and light-sheet microscopes has revolutionized the ability of scientists to visualize biological specimens (*5–8*).

One particularly useful application of deep-tissue volumetric imaging is the ability to fully appreciate 3D cellular and tissue morphology, which bypasses the need to section through different angles/axes to piece together tissue morphology. By segmenting individual cells, scientists can now quantify individual 3D cell properties such as volume, sphericity, and orientation instead of area, roundness, and other incomplete 2D cell properties obtained from often oblique tissue slices. In addition, 3D images also reveal morphological characteristics, such as protrusions and other cell-cell or cell-matrix interactions that would otherwise be lost. Recent advances in image processing techniques driven by a computer’s graphics processing unit (GPU) have further improved our understanding of cell biology by accurately segmenting both individual nuclei and entire cell bodies in an automated fashion with little-to-no user input (*9, 10*). Such tools now allow one to track the 3D morphology of thousands of cells at once.

We find that in order to image entire cleared tissue samples and to automatically segment each individual cell within the tissues, several requirements must be met: A) a high numerical aperture (N.A. ≥ 0.8), high magnification (≥ 25×) microscope objective is needed to obtain the necessary resolution to properly segment cell boundaries; B) the objective must have a long working distance, or enough to image through the entire tissue, ideally ≥ 1 mm; C) the objective, tissue, and tissue mounting medium should all have the same refractive index to produce a homogeneous immersion system, where the RI of tissues is typically ≈ 1.45 – 1.5; D) the cells must be stained with an appropriate marker that delineates cell boundaries. Unfortunately, light-sheet microscopy does not offer the resolving power needed to segment individual cells, and with the exception of a single Leica FLUOTAR objective (25x/1.0, WD 6 mm, multi-immersion) there are currently no confocal objectives on the market that fulfill criteria A-C: higher power oil and glycerol objectives are limited to short working distances, typically < 200 μm; and longer working distance, high RI objectives are limited to lower magnification (< 20×) and typically designed for light sheet microscopes. Finally, segmenting cell boundaries by either cytoplasmic or cell membrane markers can be challenging. While transgenic model species, such as mice and zebrafish, can be engineered to express genetically encoded membrane-targeted fluorescent proteins for imaging, cell boundaries in wild-type animals and non-model species must be labeled exogenously. To this end, we and others have observed that lipophilic fluorescent dyes, such as DiI, and markers of filamentous actin (F-actin), such as phalloidin, are incompatible with existing clearing methods as lipids and/or staining signals are lost during the clearing process (*11*).

To overcome these limitations, we employ a novel combination of techniques. First, since we find phalloidin to be the most consistent marker for cell body segmentation across various animal species, we devise a method to retain its signal during the clearing process. Next, we take advantage of high power, long working distance, water dipping objectives commonly available from all the major confocal objective manufacturers (Zeiss, Leica, Nikon, Olympus) by digesting the tissues with proteinase K, which renders them optically transparent in water. To prevent the enzymatic digestion from completely dissolving structures of interest, the antibodies and other fluorescent proteins are covalently bound into polyacrylamide network using a heterobifunctional linker (*12*). As a proof of principle, we use the developing mouse and catshark embryos as model systems, focusing on oral dental lamina and dermal denticle structures. Our results show that the described technique allows us to fully segment each individual cell in the tissues of interest and quantify their 3D morphological characteristics with both speed and precision.

## Results and Discussion

### Refractive index matching solutions improves optical penetration depth but disrupts fluorescent phalloidin labeling

To fully appreciate how tissues develop, it is desirable to fully characterize individual cell morphologies in their native 3D form. Cell body segmentation, however, requires a detectable marker that delineates cell boundaries; in mice, this is generally achieved using genetically encoded fluorescent proteins that label the cell membrane or nucleus. For example, *K14*^*Cre*^;*R26*^*mT/mG*^ mice express a *Keratin 14* promoter-driven Cre recombinase that activates the expression of membrane GFP in ectodermally-derived epithelial tissues, such as the developing dental epithelium. In mesenchymal cells where the Cre is inactive, membrane tdTomato is expressed. These fluorescent signals can thus be used to delineate both individual epithelial and mesenchymal cells.

A key challenge in visualizing cell boundaries at deep penetration depths is inherent tissue opacity. For instance, when embryonic day (E) 12.5 *K14*^*Cre*^;*R26*^*mT/mG*^ mouse mandibles, counterstained with DAPI and phalloidin, were imaged with a confocal microscope using a 40×/1.0 water dipping objective, the entire incisor tooth bud could be visualized, however, resolution of cells was severely limited beyond ∼100 µm (Figure 1A). To overcome this limitation, mouse mandibles were simply incubated in a refractive index matching solution, CUBIC-R (*13*), rendering the tissue optically transparent. Using a 25×/0.8 glycerol immersion objective, we were able to visualize cellular boundaries and tissue structures deep into the mesenchyme, beyond the incisor tooth bud (Figure 1B-D, Video 1).

**Figure 1.**
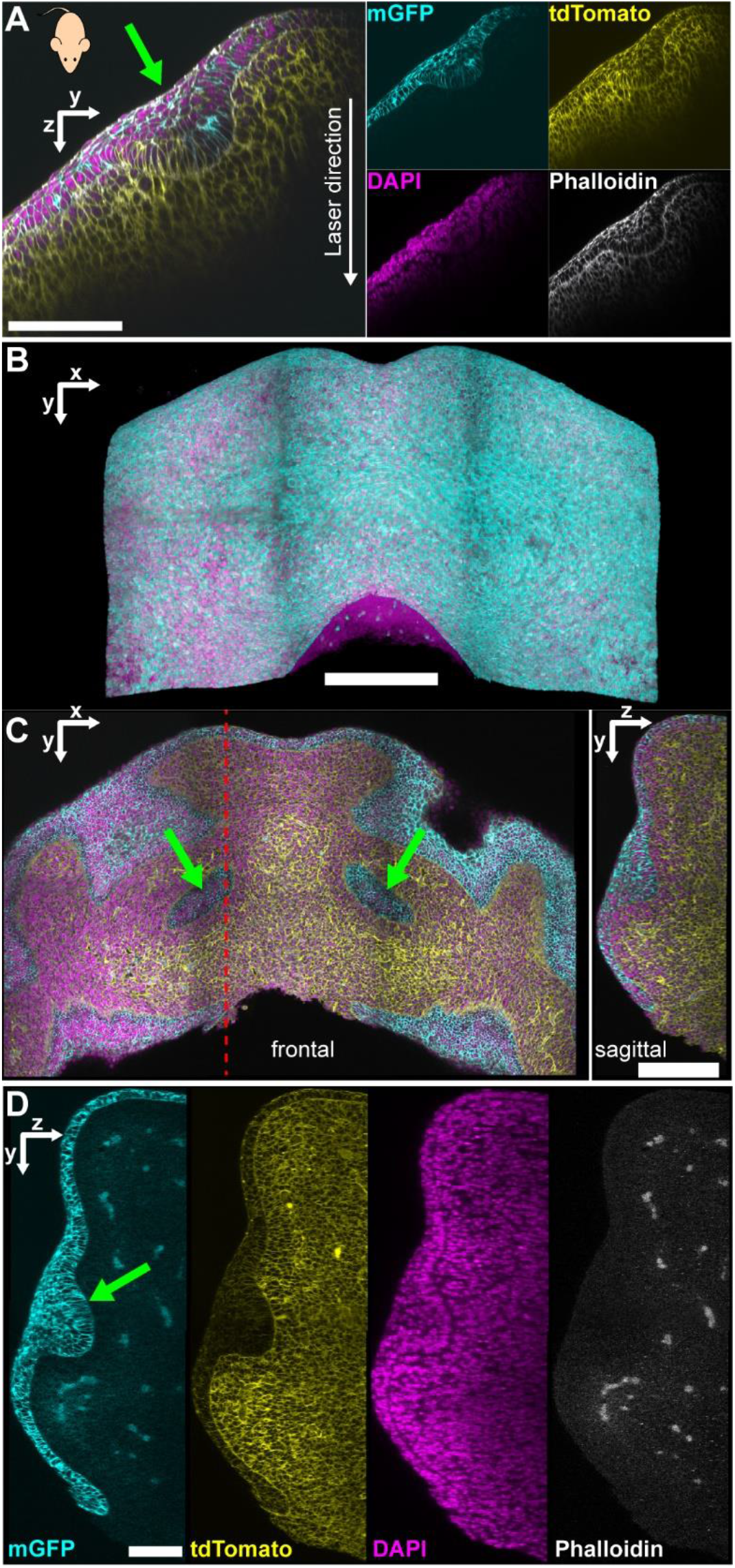
Example confocal images of E12.5 *K14*^*Cre*^;*R26*^*mT/mG*^ mouse mandibles counterstained with DAPI and phalloidin-Alexa Fluor 633. Epithelial cells are labelled by membrane GFP (mGFP), mesenchymal cells retain expression of membrane tdTomato, while all cells are labelled by phalloidin and DAPI. Green arrows indicate location of the incisor tooth buds. A) Virtual XZ-plane section of a 3D confocal volume without any tissue clearing using a 40×/1.0 water dipping objective. Image shows a sagittal view of a E12.5 *K14*^*Cre*^;*R26*^*mT/mG*^ mouse mandibular incisor tooth bud. B-D) Example 3D confocal volume cleared with CUBIC-R refractive index matching media. Image acquired using a 25× glycerol immersion objective. B) Reconstructed 3D image of the mandible (dorsal view). C) Virtual transverse (left) and sagittal (right) sections of the 3D confocal volume showing the incisor tooth bud. Dashed red line indicates the location of the sagittal plane through the tooth bud. D) Single channel images of the sagittal section. Note that phalloidin stain is not maintained after tissue clearing. Scalebars: A) 100 µm B) 300 µm; C) 200 µm; D) 100 µm.

While signals from genetically encoded fluorescence (membrane GFP and tdTomato) and DAPI were preserved after RI-matching, phalloidin staining was lost (Figure 1D). Ultimately, we were unable to find any established refractive index matching agents that did not disrupt phalloidin binding. This is unfortunate since actin is highly conserved, phalloidin exhibits broad species reactivity (14), and serves as a good marker of cell boundaries (Figure 1A). We observed strong colocalization between tdTomato and phalloidin signals in E11.5 *K14*^*Cre*^;*R26*^*mT/mG*^ mouse mandibles (Figure S1A,B), and cell segmentation results based on these two markers produced highly similar cell boundary delineations. (Figure S1C).

### Fluorescent phalloidin labeling is maintained after tissue clearing using anti-fluorophore antibodies

As proof-of-principle and to provide a deeply divergent comparison to the embryonic mouse, we utilized catshark embryos, a non-traditional model species, to perform whole-tissue imaging of cellular boundaries. Our goal was to develop a protocol applicable to a wide variety of animal models. For wild-type or non-traditional model animals that do not express fluorescent proteins labelling cell membranes, we sought alternative methods to fluorescently tag cell boundaries.

Given that phalloidin proved to be an effective marker for cell boundaries in mice, and considering the broad compatibility of tissue clearing protocols with antibody staining, we evaluated the efficacy of anti-pan actin antibodies for labeling cell boundaries. While the anti-pan actin antibodies were reactive with catshark tissues, they did not label cell boundaries as effectively as phalloidin (Supplementary Figure 2). Specifically, the antibodies failed to adequately label apical epithelial cells and certain mesenchymal cells. We next tested a panel of potential plasma membrane markers, including plasma membrane calcium-transporting ATPase 1(PMCA1); sodium potassium ATPase (Na/K ATPase); and pan cadherin (Figure S3). Although both anti-PMCA1 (Figure S3A) and pan-cadherin (Figure S3C) antibodies were reactive with catshark tissues, they only successfully labeled epithelial cell boundaries, and not the mesenchyme. Sections stained with either mouse (Figure S3B) or rabbit (Figure S3D) anti-Na/K ATPase antibodies showed no specific binding in either epithelial or mesenchymal cells.

Next, we examined non-antibody-based cell membrane markers. Wheat germ agglutinin (WGA), a lectin that binds to glycoconjugates, when conjugated to fluorescent dyes is used to label cell membranes. Additionally, lipophilic dyes, such as DiI and its derivatives, are also used to stain lipid bilayers. Thin sections of catshark mandibles were stained with WGA or the DiL derivative, CellBrite, with phalloidin stained sections used as a control (Figure S4). CellBrite successfully labeled cell boundaries (Figure S4A); however, its labeling was severely disrupted when exposed to 0.1% Triton-X (Figure S4B) or CUBIC-R (Figure S4C). A notable advantage of WGA, a protein, is that it contains primary amines and can be crosslinked into place using formaldehyde, ensuring signal stability during optical clearing (Figure S4E, F). We found, however, that WGA acted more as a broad stain for non-nuclear cell matter, and its labeling of cell boundaries was less precise compared to phalloidin (Figure S4D).

Given that phalloidin labeling was identified to be an accurate marker for cell boundaries (Figure S1), but the signal was lost after RI-matching (Figure 1D), we sought methods to retain the phalloidin signal following optical tissue clearing. Inspired by previously reported techniques that stabilize phalloidin binding (*11*), we developed a method to preserve phalloidin as a cell boundary marker for high-resolution whole-tissue imaging (Figure 2). In our approach, phalloidin-Alexa Fluor 488 was labeled with an anti-Alexa Fluor 488 antibody (Figure 2A, B). After a post-antibody labeling fixation step, the tissues were incubated in a refractive index matching solution (Figure 2C), allowing for optical clearing without disrupting phalloidin binding patterns (Figure S5E, F). Additionally, we confirmed that anti-Alexa Fluor 488 alone did not interfere with phalloidin binding (Figure S5B, C). Although Alexa Fluor 488 secondary antibodies significantly enhanced the phalloidin signal (Figure S5E), better results were achieved using a non-488 nm secondary antibody (Figure S5F). When applied to whole-mount embryonic catshark tissues, this technique preserved the phalloidin signal after refractive index matching, enabling imaging at sub-micron resolutions at full tissue penetration using confocal microscopy (Figure 3A).

**Figure 2.**
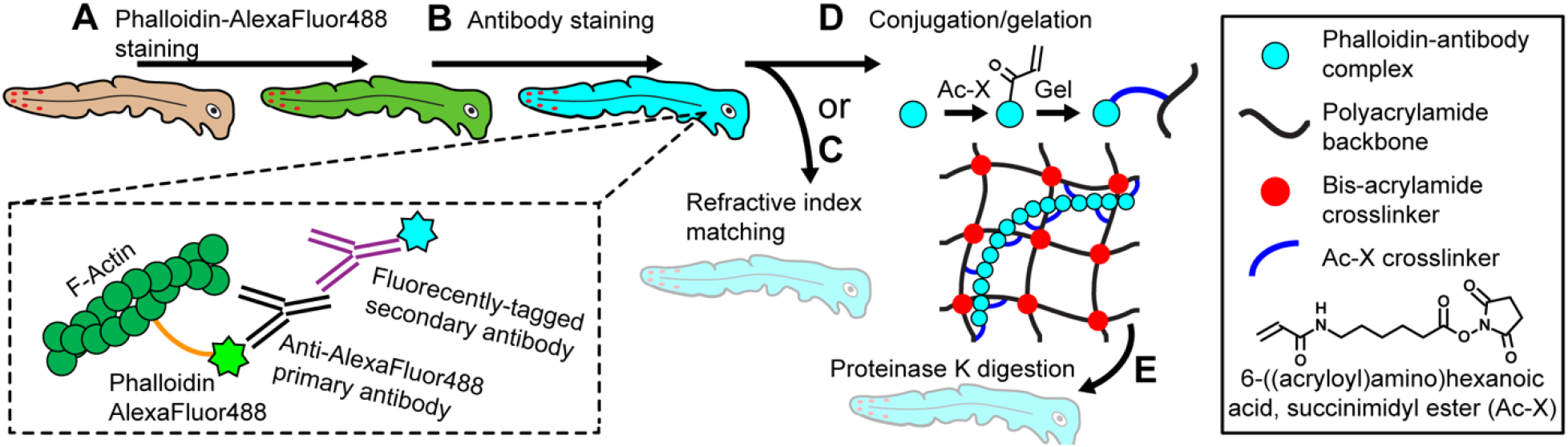
Schematic of phalloidin retention and tissue clearing process using catshark embryos as an example. A) After fixation, permeabilization, and blocking, tissues are stained with phalloidin conjugated to Alexa Fluor 488. B) Phalloidin is then antibody-tagged with anti-fluorescent dye (Alexa Fluor 488) primary antibodies, followed by fluorescently-tagged secondary antibodies. C) At this point, the sample can undergo refractive index-matching imaging. D) Alternatively, tissues can be incubated in heterobifunctional linker (Ac-X) that converts primary amines on the antibodies to reactive acrylamide groups. These acrylamide groups are then polymerized along with acrylamide monomers and bisacrylamide crosslinkers into a polyacrylamide hydrogel network. E) From here, the tissue can be digested with proteinase K to RI-match the tissue to water.

**Figure 3.**
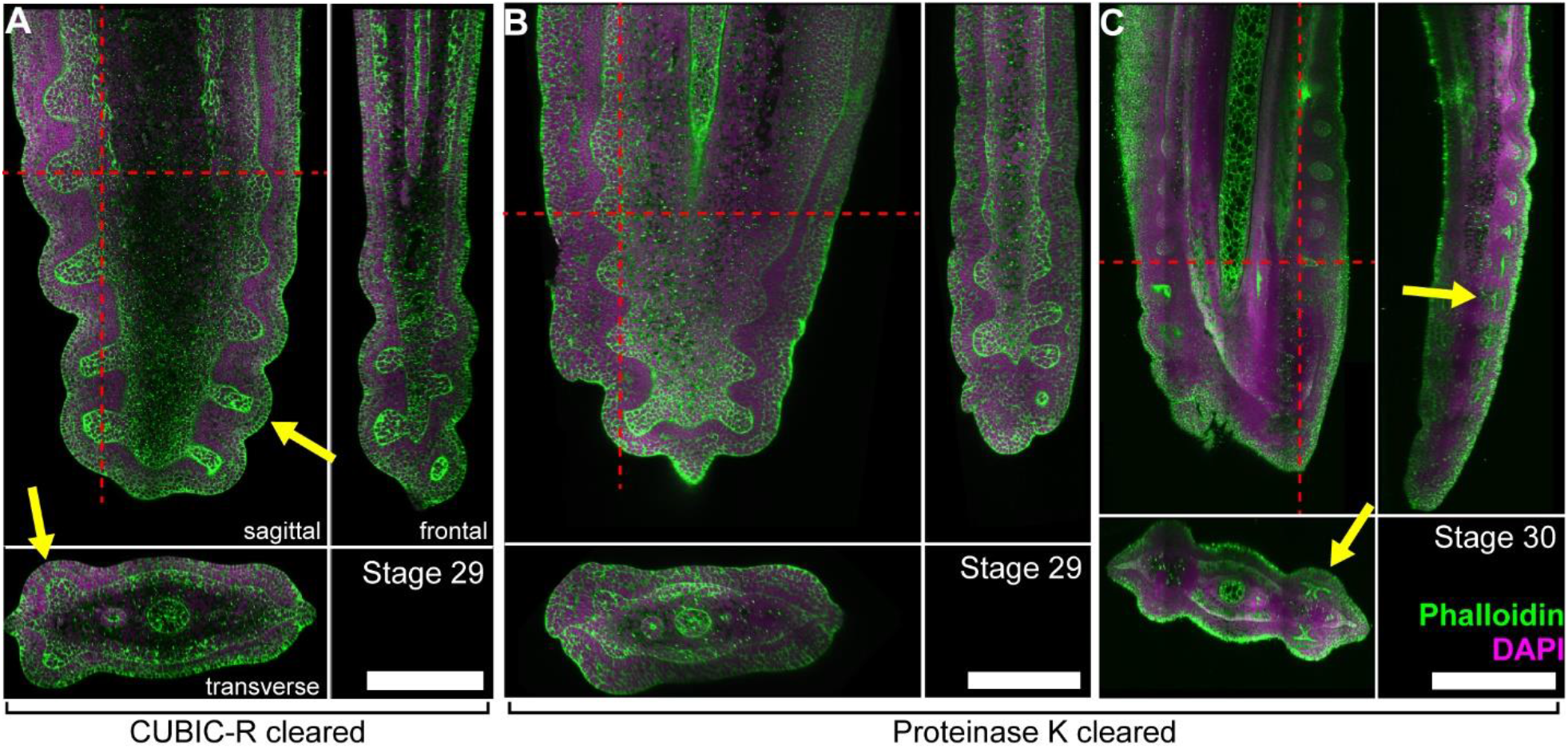
Orthogonal views of 3D confocal images catshark tail tips showing the dermal denticles (yellow arrows). Red lines indicate the location of the orthogonal sections. Tissues cleared using proteinase K or CUBIC-R, with phalloidin retention protocols applied and counterstained with DAPI. A) 27 mm stage 29 B) 27 mm stage 29 and C) 32 mm stage 30 catshark tails. Scalebars: A, B) 300 µm; C) 600 µm;

### Whole tissue imaging is accomplished using proteolytic digestion and long working distance water dipping objectives

A major limitation of the protocol outlined in Figure 2C is that high powered(≥ 25×) glycerin or oil immersion objectives are typically manufactured with limited working distances (≤ 570 µm for Zeiss, Olympus, and Nikon objectives and all but one Leica objective). This restricts full-thickness imaging of larger tissue samples (Figure S6). To address this limitation, we sought a method enabling the use of high-magnification water-immersion objectives, which provide longer working distances (> 1 mm) and are widely available across major confocal objective manufacturers. Consequently, we adapted the protein-retention Expansion Microscopy (proExM) technique (*6, 12, 14*). After antibody labeling, we conjugated the tissue proteins and antibodies to a polyacrylamide hydrogel (Figure 2D) and subjected the tissue-gel construct to nonspecific proteolytic digestion. This process rendered the samples transparent in water, and compatible with water immersion objectives (Figure 2E).

Protein-gel conjugation was achieved using a heterobifunctional linker that covalently substitutes primary amines of tissue protein and antibodies with reactive acrylamide groups, which subsequently polymerize into the backbone of the polyacrylamide gel. Using a water immersion objective, we successfully imaged catshark tail-tip tissues, which are several hundred microns thick, at high resolution and resolving power while preserving the phalloidin signal (Figure 3B,C, Video 2). The quality of the protein-retention, gel embedded images (Figure 3B) was comparable to that of similarly sized samples cleared using CUBIC-R (Figure 3A). Moreover, the gel embedded samples allowed for imaging with higher-power objectives, significantly enhancing the potential resolution, especially in the z-direction.

To further evaluate the performance of the protein-retention, gel embedding technique in other tissues and developmental stages, we imaged a series of catshark tail tips (Figure 4A-D), and mandibles (Figure 4E-G). Special attention was given to comparing the dermal denticles of the tail tip and the mandibular dental lamina and tooth buds. Additionally, we applied this technique to E12.5 *K14*^*Cre*^;*R26*^*mT/mG*^ mouse mandibles, where fluorescent proteins were genetically encoded (Figure S7). In all instances, we were able to obtain detailed 3D images of the cell boundaries.

**Figure 4.**
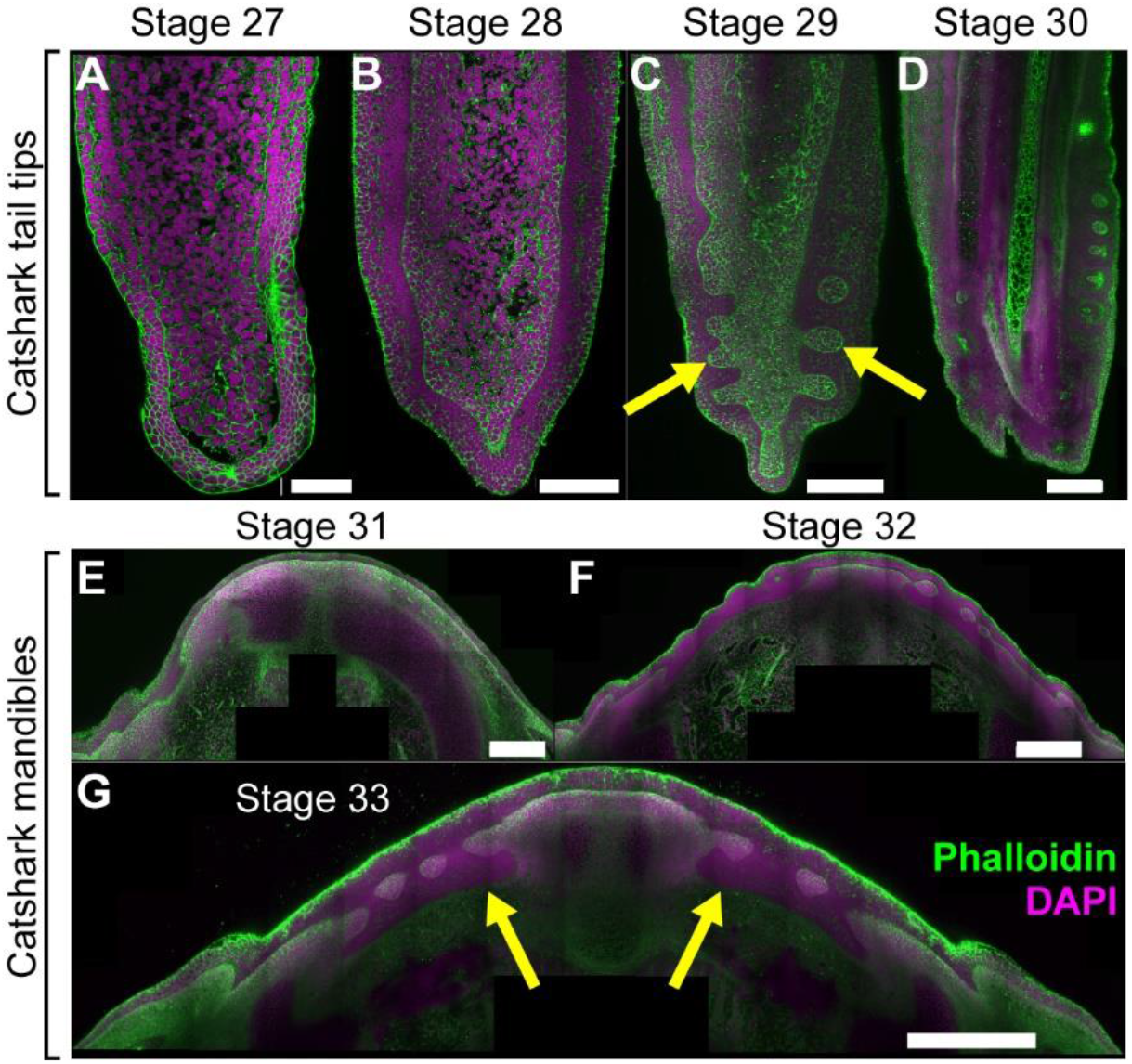
3D confocal images of catshark tissues cleared using proteinase K. A-D) Example virtual sagittal slices from the confocal volumes through the tail tips showing the dermal denticles (yellow arrows): A) 24 mm stage 27 shark, B) 25 mm stage 28 shark, C) 27 mm stage 29 shark, and D) 32 mm stage 30 shark. E-G) Virtual frontal slices through the mandibles showing the dental lamina (yellow arrows): E) 43 mm stage 31 shark mandible, F) 48 mm stage 32 shark mandible, and G) 50 mm stage 33 shark mandible. Scalebars: A) 80 µm; B) 150 µm; C) 250 µm; D) 300 µm; E) 300 µm; F) 400 µm; G) 500 µm.

### Imaging of optically cleared tissues with intact cell boundary markers enables full tissue cell segmentation

We next evaluated our ability to segment individual cells following whole-tissue confocal imaging. Using the tail tip from a 24 mm (stage 27) catshark as an example, we automatically segmented each cell within the tissue using a custom-trained deep-learning segmentation model (*15*). The phalloidin channel (Figure 5A), highlighting cell boundaries, provided the primary input for the model, allowing for successful segmentation of all cells (Figure 5B, Video 3). These segmentations were then used to produce 3D heat maps quantifying various individual cell properties such as: average F-actin concentration per cell (Figure 5C); the length of the longest principle axis of the cell (Figure 5D); cell sphericity (Figure 5E); the ratio of major to minor principal axes (Figure 5F); the local cell density, in units cells/unit volume (Figure 5G); and individual cell volume (Figure 5H).

**Figure 5.**
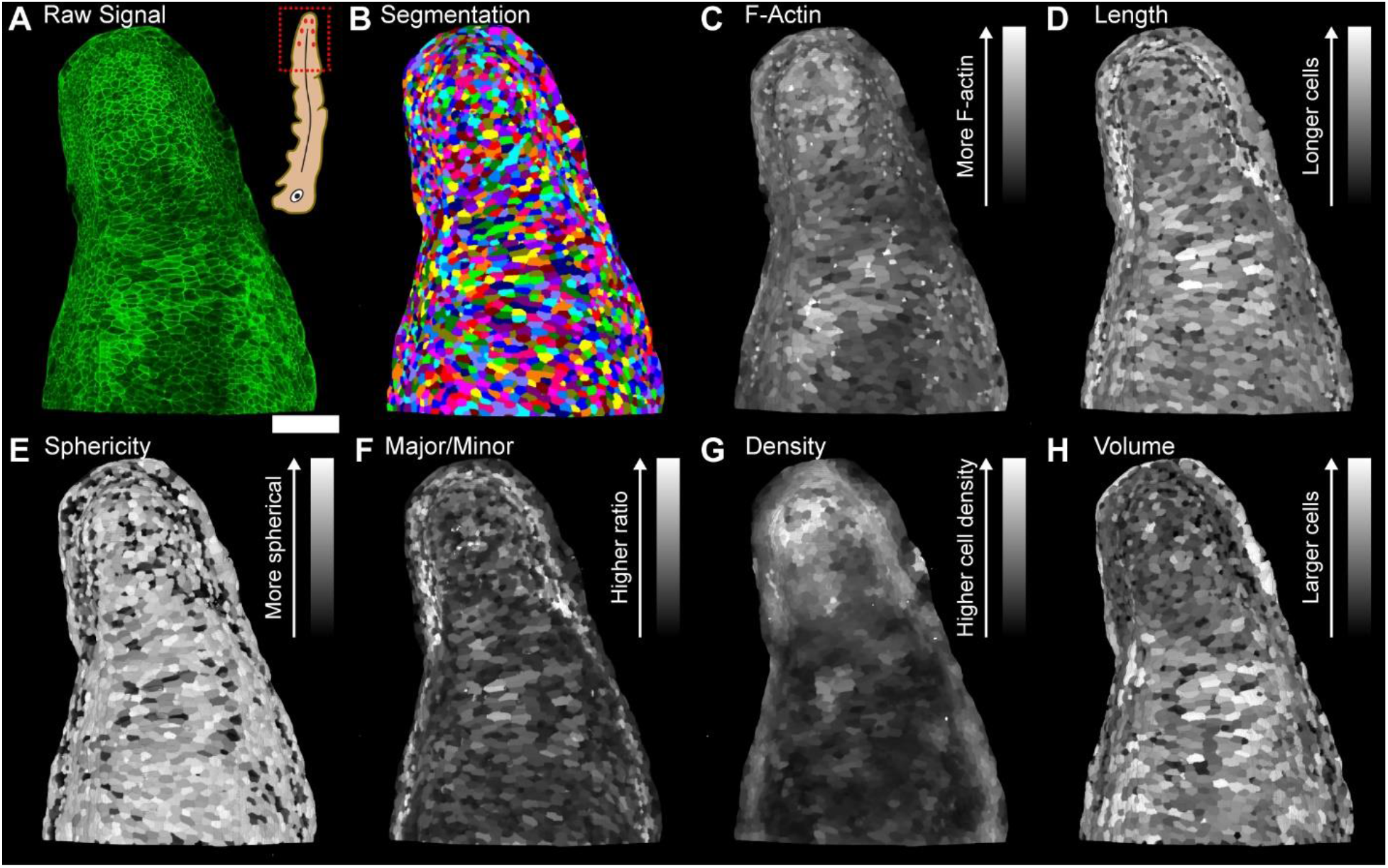
Example tail tissue from a 24 mm (stage 27) shark cleared and imaged to obtain individual 3D volumetric cell morphology properties. Volumetric images of A) input phalloidin fluorescence signal, and B) individually segmented cells. C-H) Heat maps of phalloidin intensity and cellular morphology. C) Average phalloidin intensity per cell, D) length of longest principal axis of each cell, E) cell sphericity, F) ratio of the largest/smallest principal axes, G) local cell density in units of cells/volume, and H) cell volume. Scalebar 100 µm.

### Heat maps of 3D cellular properties to better understand epithelial bending modes

Epithelial morphogenic processes such as bending of epithelial sheets, in simple terms, can occur through two different modes: invagination into the underlying tissue; or evagination outwards from the epithelial surface (*16*). The mechanisms by which epithelial bending direction is determined, however, are not well understood. Using the catshark mandibular dental lamina and tail-tip dermal denticles as examples, we compared the cellular properties of both invaginating (dental lamina) and evaginating (dermal denticles) tissue structures (Figure 6A). To analyze cellular morphology in specific regions and orientations within the tissue interior, we generated virtual sections (Figure 6B) through the 3D cell property heat maps, providing 2D snapshots of the 3D cell properties (Figure 6C-F). To explore the cell behaviors that could lead to these two different types of epithelial bending, we examined the length of the longest principal axis of the cells (Figure 6C), the cell sphericity (Figure 6D), the cellular volume (Figure 6E), and the distribution of F-actin across the tissues (Figure 6F).

**Figure 6.**
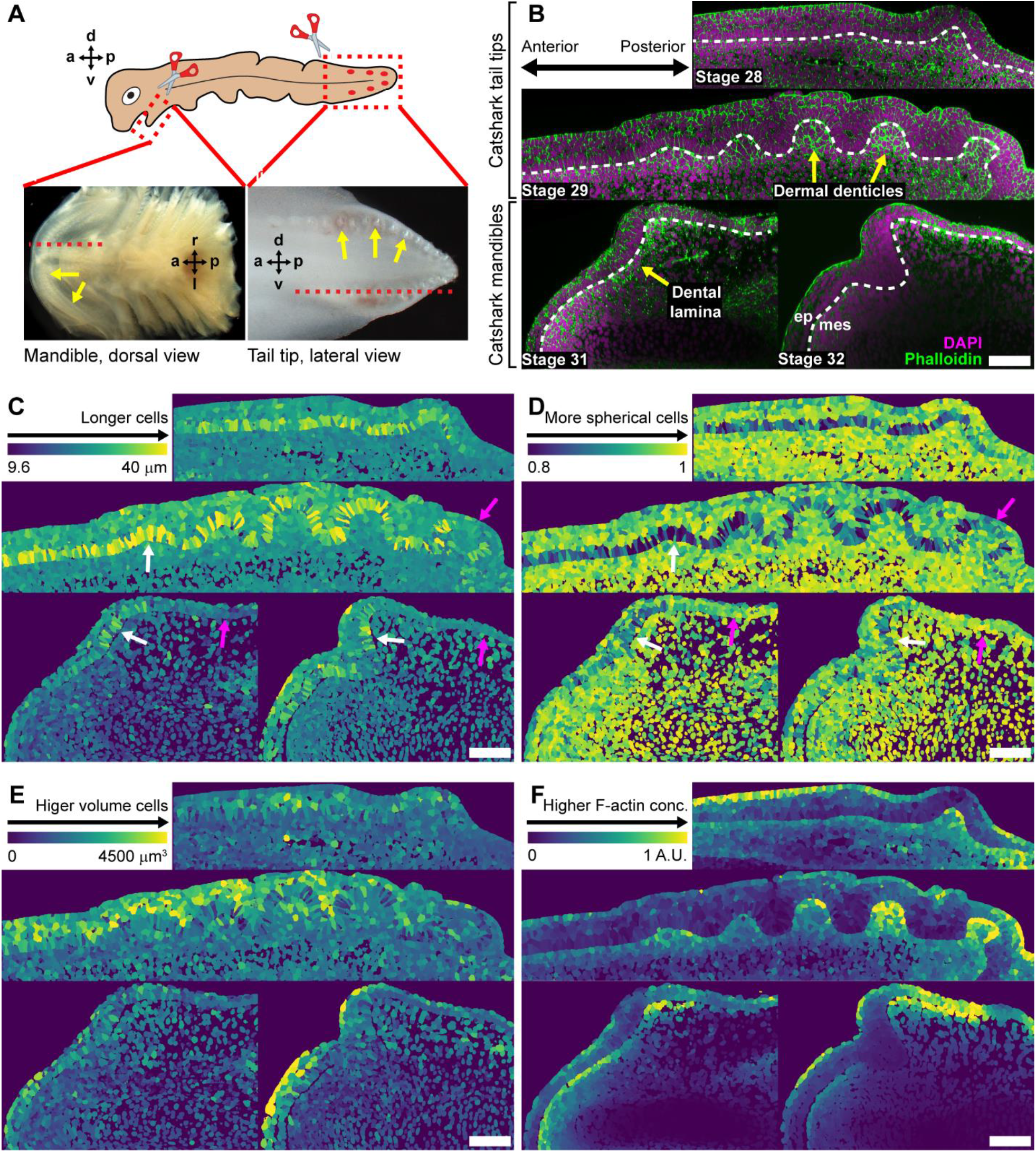
Heat maps of 3D cell properties mapped on a cell-by-cell basis from 3D volumetric images. A) Diagram of catshark i) mandible showing the odontogenic band (yellow arrows) and ii) tail tip showing the dermal denticles (yellow arrows). Dorsal (d), ventral (v), anterior (a), posterior (p), left (l), and right (r) anatomical directions. Red lines indicate location of the virtual sections displayed in panels B-F. B) Representative sections of 25 mm stage 28 shark tail, 28mm stage 29 shark tail, 43 mm stage 31 shark mandible, and 47 mm stage 32 shark mandible stained with phalloidin and DAPI. Yellow arrows indicate the dermal denticles and dental lamina of the tail and mandible, respectively. White lines indicate the boundary between the epithelium (ep) and mesenchyme (mes). Cell-by-cell measurements of: C) length of the longest principal axis of the cells, white and magenta arrows indicate regions with longer and shorter basal epithelial cells, respectively; D) cell sphericity (unitless), white and magenta arrows indicate regions with more elliptical and more spherical basal epithelial cells, respectively; E) cell volume; and F) average cellular F-actin staining intensity. Scalebars 100 µm.

While the first signs of tooth development and invagination occur when the local epithelium thickens (*17*) and the basal epithelial cells elongate (*18*), it is less clear if basal epithelial cell elongation also precedes epithelial evagination in dermal denticles. We first confirmed that the basal epithelial cells of the catshark mandibular dental lamina are elongated (Figure 6C, white arrows) than those more oral and aboral (magenta arrows) in both early (stage 31) and later (stage 32) dental lamina. Cell sphericity is a similar metric that measures how spherical or elliptical a cell is; thus, we expect that basal epithelial cells at the dental placode to be more elliptical. Indeed, we find that the same cells that are elongated are also more elliptical (Figure 6D). Next, we find that similar to the dental placode, the basal epithelial cells of the evaginating tail-tip dermal denticles also locally elongate (Figure 6C) and become more elliptical (Figure 6D) in regions of placode formation in both early (stage 28) and later stage (stage 29) denticles. Interestingly, the basal epithelial cells of newly formed dermal denticle placodes are more elongated and elliptical (white arrows) than older, more distal placodes that have undergone greater evagination (magenta arrows). Next, we examined whether cell growth, by means of observing local changes in cell volume, predicted epithelial bending. Surprisingly, we find no definitive patterns of cell volume in either the catshark dental lamina or dermal denticles (Figure 6E). Finally, we look at the distribution of F-actin staining intensity on a cell-by-cell basis by averaging the staining intensity through the whole volume of each cell (Figure 6F). Mesenchymal cells underlying the dermal denticles exhibit elevated F-actin staining intensity that indicates elevated cell tension. In contrast, we see no particular pattern of F-actin staining intensity in the dental lamina placodes, except for a potential loss of F-actin in the mesenchyme directly beneath the dental lamina. Together, these experiments demonstrated that our clearing protocol is suitable for studying physical features of cells and indicated that these changes may underlie epithelial morphogenetic events.

## Conclusions

In this work we developed a method to image cell boundaries at sub-micron resolutions of entire tissues. While genetic model organisms can be bred to express fluorescent proteins in their cell membranes, wild type and non-traditional model animals do not have this luxury. Given that phalloidin, which selectively labels F-actin in most species, acts as a good marker of cell boundaries, we adapted a method to retain phalloidin staining during tissue clearing. These phalloidin-retained tissues were further processed and optically cleared by two different methods: the embryonic tissues could simply be incubated in a refractive index-matching solution; or embedded into a polyacrylamide gel, covalently binding the tissue proteins to the gel backbone, and digesting the remaining tissue so that tissue was refractive index-matched with water. The latter method allowed us to utilize high magnification, long working distance water dipping objectives to image the entire tissue. The subsequent images were then automatically segmented using a user-trained neural network segmentation model, which output each individual cell volume of the tissues. As an example application, we examined epithelial bending of the invaginating catshark dental lamina and their evaginating dermal denticles. These data suggest that basal epithelial cells elongate in regions of high in-plane epithelial compression, and that increased F-actin activity in the mesenchymal portion of the dermal denticle placode constricts the mesenchyme, causing it to buckle outwards towards the epithelium, driving its evagination. We expect that the techniques devised in this work, will help developmental biologists apply more quantitative measures of 3D cell morphology to understand tissue morphogenesis.

## Materials & Methods

### Materials

Tricaine (Syndel), Phosphate Buffered Saline (PBS) (Fisher Scientific), paraformaldehyde (PFA, Thermo Scientific), goat serum (Gibco), bovine serum albumin (BSA, Fisher Scientific), Triton X-100 (Fisher Scientific), phalloidin conjugated to either Alexa Fluor™ 488 (Thermo Fisher A12379), or Alexa Fluor™ 633 (Thermo Fisher A22284), 4′,6-diamidino-2-phenylindole (DAPI, Biotium 40009), rabbit anti-Alexa Fluor™ 488 (Thermo Fisher A11094), 35 mm glass bottom dish (Thermo Scientific), low melting point agarose (Thermo Scientific), #1.5 coverglass (Fisher Scientific), dimethyl sulfoxide (DMSO, Thermo Scientific), acrylamide (AAm, Thermo Scientific), bis-acrylamide (BAAm, Thermo Scientific), ammonium persulfate (APS, Acros Organics), tetramethylethylenediamine (TEMED, Thermo Scientific), 4-hydroxy-2,2,6,6-tetramethylpiperidin-1-oxyl (TEMPO, Thermo Scientific), and Proteinase K (Invitrogen) were used as purchased.

A wash solution of 0.1% Triton X in 1× PBS (PBST), CUBIC-R refractive-index matching solution (45 w/v% antipyrine (Thermo Scientific), 30 w/v% nicotinamide (Thermo Scientific), and 0.5 v/v% N-butyldiethanolamine (Thermo Scientific) in distilled water as described previously (*13*), were made fresh in the lab. Digestion buffer consisting of Triton X, ethylenediaminetetraacetic acid (EDTA, Sigma), tris(hydroxymethyl)aminomethane (Tris hydrochloride, Roche), and sodium chloride (Sigma) was prepared as previously described (*12*).

6-((Acryloyl)amino)hexanoic Acid, Succinimidyl Ester (AcX, CAS number: 63392-86-9) was synthesized as previously described (*19, 20*).

### Collection of embryos and fixation

#### Mice

*K14*^*Cre*^;*R26*^*mT/mG*^ mice were generated by crossing *K14*^*Cre*^ (MGI:2445832) (*21*) with *R26*^*mT/mG*^ (MGI:3716464) (*22*). C57BL/6 were purchased from JAX. All mice were group housed and genotyped as previously published. To generate embryos for experiments, mice were mated overnight, and noon of the day of vaginal plug discovery was designated as E0.5. Pregnant females were euthanized by CO_2_ followed by cervical dislocation. Both male and female GFP-positive *K14*^*Cre*^ ;*R26*^*mT/mG*^ embryos or C57BL/6 embryos were selected at random and used for experiments. Mandibles were dissected out in PBS and fixed in freshly made 4% paraformaldehyde (PFA) in 1× PBS for 12 hours on a rocker at 4°C. The samples were then washed in 1× PBS three times for 10 minutes each and stored in 0.5% PFA at 4°C until use. All mice were maintained in the University of California Los Angeles (UCLA) pathogen-free animal facility. All animal procedures were conducted in compliance with animal protocols approved by the UCLA Institutional Animal Care and Use Committee (Protocol Number ARC-2019-013).

#### Sharks

Fertilized eggs of small spotted catshark (*Scyliorhinus rotifer*) were obtained from the Marine Biological Laboratory, Woods Hole, MA (stages 28-32). Embryos were mechanically extracted and placed in a 0.5% Tricaine solution diluted in in 1× PBS solution for thirty minutes. Embryos were then fixed in freshly made 4% PFA in 1× PBS for 24 hours on a rocker at 4° C. Fixed samples were washed in 1× PBS four times for 1h each. Mandibles and tails were dissected out and stored in 1× PBS. From this point forward, samples were protected from light during.

### Immunofluorescence labeling

#### Thin sectioning

Fixed tissues were stepped into 30% sucrose before embedding in OCT and cryosectioning (Thermo Fisher NX50) 10 µm thick sections. Sections were dried and stored at -80°C. Sections were rehydrated in PBS before use.

A. Antibody and fluorescence labeling. Sections were incubated in blocking buffer (5% goat serum, 1% BSA, 0.3% Triton X in 1× PBS) for 1 hour. The sections were then incubated in primary antibodies overnight at 4°C: monoclonal Rabbit anti-PMCA1 (1:200, abcam, ab190355); monoclonal mouse anti-alpha 1 sodium potassium ATPase antibody (1:200, abcam, ab7671); monoclonal rabbit anti-pan cadherin antibody, intercellular junction Marker (1:200, abcam, ab51034); and monoclonal rabbit anti-alpha 1sodium potassium ATPase antibody (1:200, abacm, ab76020) in blocking buffer. Sections labeled with anti-pan actin (1:300, Cytoskeleton, AAN02) were labeled according to the manufacturer’s protocol. The next day, the sections were washed with PBS and incubated in secondary antibodies (1:300, goat anti-rabbit 633, Biotium 20122) and DAPI (1:500) in blocking buffer for hours. Sections were then washed with PBS and mounted with coverglass.
B. Membrane labeling. To test different strategies to label cell membranes, thin sections were labeled with either a DiI-based lipophilic dye (CellBrite Orange, Biotium 30022) or fluorescent wheat germ agglutinin (WGA CF555, Biotium 29076). Sections stained with CellBrite were permeabilized with 0.1% Triton-X in PBS for 10 minutes (if noted), washed with PBS, incubated in CellBrite (1:300 in PBS) for 10 minutes, washed with PBS and mounted in CUBIC-R or PBS. Sections stained with WGA were incubated in WGA (1:200 in PBs) for 30 minutes, washed with PBS, fixed in 1% PFA in PBS for 20 minutes, permeabilized with 0.1% Triton-X in PBS for 30 minutes, washed with PBS and mounted in PBS or CUBIC-R.
C. Anti-Alexa Fluor™ 488 labeling. To test the ability to retain phalloidin labeling using thin sections, sections were incubated in blocking buffer (5% goat serum, 1% BSA, 0.3% Triton X in 1× PBS) for 1 hour followed by phalloidin-Alexa Fluor™ 488 (1:200, ThermoFisher, A11094) in blocking buffer overnight at 4°C. Sections were washed with PBST. At this point, positive controls were mounted in PBS, and negative controls were mounted in CUBIC-R with no further primary or secondary antibodies. Sections were then incubated in anti-fluorophore antibody (1:200) in blocking buffer overnight at 4°C. Samples without secondary antibody labeling were mounted in PBS or CUBIC-R at this point. Sections were then incubated in secondary antibodies (1:300) in blocking buffer for 2 hours. Sections were mounted in CUBIC-R.

### Whole mount antibody and fluorescence labeling

Mouse mandibles and shark mandibles and tails were first incubated in blocking buffer (5% goat serum, 1% BSA, 0.3% Triton X in 1× PBS) for 24 hours on a rocker at 4° C. Next, the samples were incubated in phalloidin conjugated to either Alexa Fluor™ 488 or 633 (1:200) and DAPI (0.1 mg/mL) in blocking buffer for 7 days on a rocker at 4° C. The samples were then washed in PBST three times for 1h each before storage in PBS. Samples receiving no further antibody treatment were either cleared or imaged directly. Samples were then stained with anti-fluorophore antibody (1:200, ThermoFisher, A11094) in blocking buffer for 3 days on a rocker at 37° C protected from light. The samples were then washed in PBST three times for 1h each. The labeled samples were then stained with secondary antibody (1:300, goat anti-rabbit 633, Biotium 20122) in blocking buffer for 3 days on a rocker at 37° C. The samples were then washed in PBST three times for 1h each.

### Tissue clearing

Tissues were rendered optically transparent by two different methods: the first renders the samples optically transparent by simply immersing them in an aqueous-based high refractive index solution; the second covalently binds the antibodies into a 3D polymer network and enzymatically digests the remaining tissue.

A. To render tissues optically transparent in a high-refractive index solution, fluorescently labeled samples were first fixed in freshly made 1% PFA in 1× PBS for 24 hours on a rocker at 4° C. Samples were then washed for 2h in 1× PBS before immersing in a for 1-3 days (depending on the size of the embryonic sample) in CUBIC-R to render the tissues optically transparent for three-dimensional imaging. Samples were mounted on a 35 mm coverglass bottom dish suspended in a 0.5% agarose solution in CUBIC-R to avoid movement of the sample during the imaging process. The mounted samples were then covered with a coverglass before imaging.
B. To render tissues optically transparent in a low refractive index aqueous solution, fluorescently labeled samples were incubated in a freshly made solution of the heterobifunctional linker AcX. A stock solution of 50 mg/mL AcX in anhydrous DMSO was freshly prepared before diluting 10-fold in PBS. Samples were immediately immersed in the AcX solution for 24 hours on a rocker at 22° C. The functionalized tissues were then washed with PBST for 2h and immersed in a cold polyacrylamide pre-polymer solution consisting of AAm (9.6 w/v%), BAAm (0.25 w/v%), TEMPO (0.75 w/v%), APS (0.2 w/v%), and TEMED (0.2 v/v%) in 1× PBS for 4h in an ice bath, while rocking. The pre-polymer solution was replaced and the samples were placed between two glass slides with 1 mm spacers, which was filled with the pre-polymer solution. The glass slide-sample sandwiches were then incubated at 37° C for 1h to polymerize the polyacrylamide gels. The tissue-gel construct was then trimmed and immersed in a solution of Proteinase K in digestion buffer (0.2 mg/mL) and incubated for 48h on a rocker at 22° C, or until the tissue was rendered optically transparent. The tissues were then washed with 1× PBS three times for 1h each. Samples were mounted in a 35 mm plastic dish suspended in a 0.5% agarose solution in 1× PBS before imaging.

### Confocal imaging

Samples were imaged with a Zeiss confocal laser scanning microscopes (LSM 900) equipped with either a Plan-Apochromat 25×/0.8 glycerine immersion objective with a working distance (WD) of 0.57 mm or a Plan-Apochromat 40×/1.0 DIC VIS-IR water dipping objective, WD 2.5mm. A pinhole of 1 AU was used. Three dimensional, whole-tissue volumes were acquired using a combination of Z-stacks and image tiling. Tissues cleared with CUBIC-R were imaged using the 25× objective and gel-embedded tissues cleared with Protienase K were imaged using the 40× objective

### Image Processing and segmentation

Images were processed and visualized with Zen (Zeiss), ImageJ and Imaris. The multi-tiled images were stitched with the ImageJ plugin BigStitcher (*23*). At times, contrast adjustments were applied to enhance details for visualization and segmentation, but not for signal quantification. The stitched images were reoriented and cropped to contain the region of interest and reduce file size and segmented using CellPose (*9*). Before segmentation the neural network model was trained using a subset of images where the phalloidin signal was used as the primary channel, and the DAPI signal was used as the secondary (*15*). Training was repeated and refined as needed to produce properly segmented 3D cell volumes. Segmented volumes were output as multi-Z masks tiff file.

Once the individual cells were segmented, the mask files were imported into MATLAB the properties of 3D volumetric image regions were measured with the regionprops3 algorithm. A piece of custom MATLAB code was then utilized to reconstruct 3D heatmaps of the segmented cells labeled with different morphological features (volume, sphericity, F-actin concentration, cell length, etc…).

## References

1. D. S. Richardson, W. Guan, K. Matsumoto, C. Pan, K. Chung, A. Ertürk, H. R. Ueda, J. W. Lichtman, Tissue clearing. Nat. Rev. Methods Prim. 1, 84 (2021).

2. D. S. Richardson, J. W. Lichtman, Clarifying Tissue Clearing. Cell. 162, 246–257 (2015).

3. T. Yu, J. Zhu, D. Li, D. Zhu, Physical and chemical mechanisms of tissue optical clearing. iScience. 24, 102178 (2021).

4. A. Azaripour, T. Lagerweij, C. Scharfbillig, A. E. Jadczak, B. Willershausen, C. J. F. Van Noorden, A survey of clearing techniques for 3D imaging of tissues with special reference to connective tissue. Prog. Histochem. Cytochem. 51, 9–23 (2016).

5. K. Chung, J. Wallace, S.-Y. Kim, S. Kalyanasundaram, A. S. Andalman, T. J. Davidson, J. J. Mirzabekov, K. A. Zalocusky, J. Mattis, A. K. Denisin, S. Pak, H. Bernstein, C. Ramakrishnan, L. Grosenick, V. Gradinaru, K. Deisseroth, Structural and molecular interrogation of intact biological systems. Nature. 497, 332–337 (2013).

6. F. Chen, P. W. Tillberg, E. S. Boyden, Expansion microscopy. Science (80-. ). 347, 543– 548 (2015).

7. E. A. Susaki, K. Tainaka, D. Perrin, F. Kishino, T. Tawara, T. M. Watanabe, C. Yokoyama, H. Onoe, M. Eguchi, S. Yamaguchi, T. Abe, H. Kiyonari, Y. Shimizu, A. Miyawaki, H. Yokota, H. R. Ueda, Whole-Brain Imaging with Single-Cell Resolution Using Chemical Cocktails and Computational Analysis. Cell. 157, 726–739 (2014).

8. N. Renier, Z. Wu, D. J. Simon, J. Yang, P. Ariel, M. Tessier-Lavigne, iDISCO: A Simple, Rapid Method to Immunolabel Large Tissue Samples for Volume Imaging. Cell. 159, 896–910 (2014).

9. C. Stringer, T. Wang, M. Michaelos, M. Pachitariu, Cellpose: a generalist algorithm for cellular segmentation. Nat. Methods. 18, 100–106 (2021).

10. S. Berg, D. Kutra, T. Kroeger, C. N. Straehle, B. X. Kausler, C. Haubold, M. Schiegg, J. Ales, T. Beier, M. Rudy, K. Eren, J. I. Cervantes, B. Xu, F. Beuttenmueller, A. Wolny, C. Zhang, U. Koethe, F. A. Hamprecht, A. Kreshuk, ilastik: interactive machine learning for (bio)image analysis. Nat. Methods. 16, 1226–1232 (2019).

11. C. E. Park, Y. Cho, I. Cho, H. Jung, B. Kim, J. H. Shin, S. Choi, S.-K. Kwon, Y. K. Hahn, J.-B. Chang, Super-Resolution Three-Dimensional Imaging of Actin Filaments in Cultured Cells and the Brain via Expansion Microscopy. ACS Nano. 14, 14999–15010 (2020).

12. S. M. Asano, R. Gao, A. T. Wassie, P. W. Tillberg, F. Chen, E. S. Boyden, Expansion Microscopy: Protocols for Imaging Proteins and RNA in Cells and Tissues. Curr. Protoc. Cell Biol. 80 (2018), doi:10.1002/cpcb.56.

13. E. A. Susaki, C. Shimizu, A. Kuno, K. Tainaka, X. Li, K. Nishi, K. Morishima, H. Ono, K. L. Ode, Y. Saeki, K. Miyamichi, K. Isa, C. Yokoyama, H. Kitaura, M. Ikemura, T. Ushiku, Y. Shimizu, T. Saito, T. C. Saido, M. Fukayama, H. Onoe, K. Touhara, T. Isa, A. Kakita, M. Shibayama, H. R. Ueda, Versatile whole-organ/body staining and imaging based on electrolyte-gel properties of biological tissues. Nat. Commun. 11, 1982 (2020).

14. P. W. Tillberg, F. Chen, K. D. Piatkevich, Y. Zhao, C. C. Yu, B. P. English, L. Gao, A. Martorell, H. J. Suk, F. Yoshida, E. M. Degennaro, D. H. Roossien, G. Gong, U. Seneviratne, S. R. Tannenbaum, R. Desimone, D. Cai, E. S. Boyden, Protein-retention expansion microscopy of cells and tissues labeled using standard fluorescent proteins and antibodies. Nat. Biotechnol. 34, 987–992 (2016).

15. M. Pachitariu, C. Stringer, Cellpose 2.0: how to train your own model. Nat. Methods. 19, 1634–1641 (2022).

16. Y. Wang, D. Stonehouse-Smith, M. T. Cobourne, J. B. A. Green, M. Seppala, Cellular mechanisms of reverse epithelial curvature in tissue morphogenesis. Front. Cell Dev. Biol. 10, 1–13 (2022).

17. J. Pispa, I. Thesleff, Mechanisms of ectodermal organogenesis. Dev. Biol. 262, 195–205 (2003).

18. J. Li, L. Chatzeli, E. Panousopoulou, A. S. Tucker, J. B. A. Green, Epithelial stratification and placode invagination are separable functions in early morphogenesis of the molar tooth. Development. 143, 670–681 (2016).

19. S. C. P. Norris, J. Soto, A. M. Kasko, S. Li, Photodegradable Polyacrylamide Gels for Dynamic Control of Cell Functions. ACS Appl. Mater. Interfaces. 13, 5929–5944 (2021).

20. D. D. Pless, Y. C. Lee, S. Roseman, R. L. Schnaar, Specific cell adhesion to immobilized glycoproteins demonstrated using new reagents for protein and glycoprotein immobilization. J. Biol. Chem. 258, 2340–9 (1983).

21. H. R. Dassule, P. Lewis, M. Bei, R. Maas, A. P. McMahon, Sonic hedgehog regulates growth and morphogenesis of the tooth. Development. 127, 4775–4785 (2000).

22. M. D. Muzumdar, B. Tasic, K. Miyamichi, L. Li, L. Luo, A global double-fluorescent Cre reporter mouse. genesis. 45, 593–605 (2007).

23. D. Hörl, F. Rojas Rusak, F. Preusser, P. Tillberg, N. Randel, R. K. Chhetri, A. Cardona, P. J. Keller, H. Harz, H. Leonhardt, M. Treier, S. Preibisch, BigStitcher: reconstructing high-resolution image datasets of cleared and expanded samples. Nat. Methods. 16, 870–874 (2019).

