## Supplementary Information for "Whole Tissue Imaging of Cellular Boundaries at Sub-Micron Resolutions for Automatic Cell Segmentation: Applications in Epithelial Bending of Ectodermal Appendages"

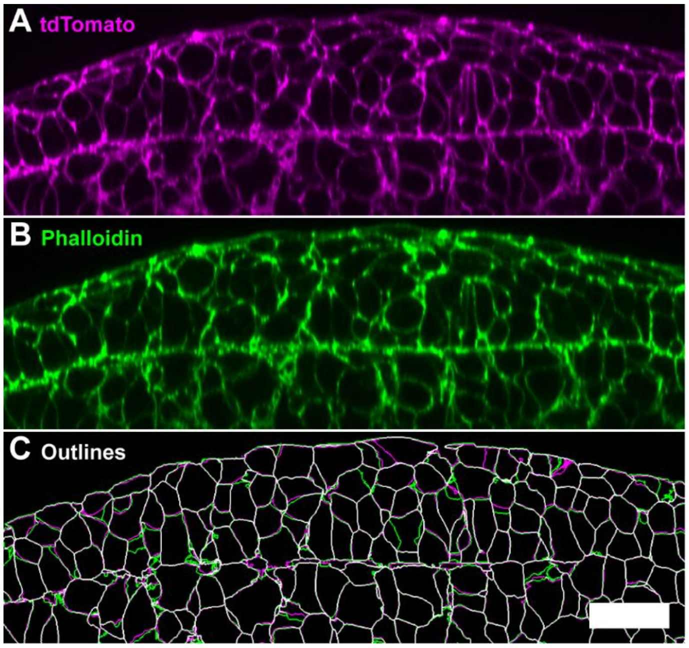


**Figure S1**. Example virtual XZ-plane section of a 3D confocal volume of a E11.5 *K14*$^{Cre}$;*R26*$^{mT/mG}$ mouse mandible counterstained with phalloidin-Alexa Fluor 633 imaged using a 40×/1.0 water immersion objective. No optical clearing agents were used. Image shows a sagittal view of mouse mandibular incisor tooth bud. A) Expression of membrane tdTomato and B) dye-conjugated phalloidin. C) Computer-generated cell boundaries segmented using CellPose software from the tdTomato (magenta lines) and phalloidin (green lines) signals. White lines indicate overlap of the cell segmentations from the two different signal types. Scalebar: 25 µm.

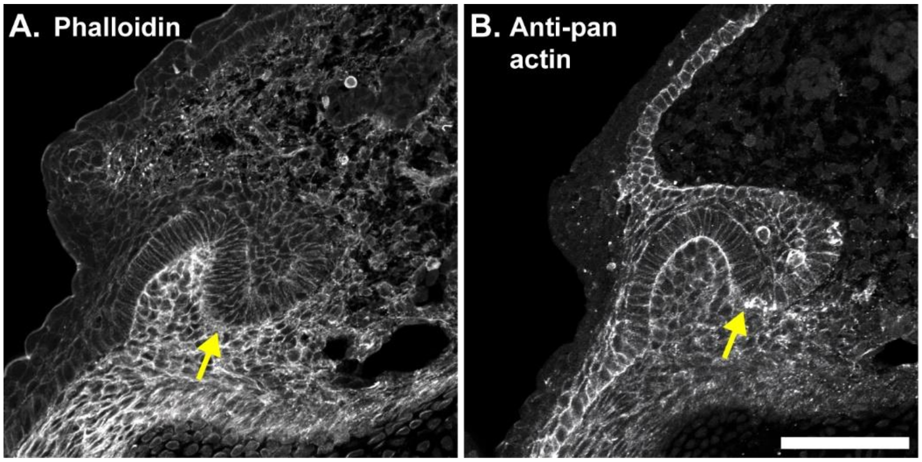


**Figure S2**. Representative serial thin sections of stage 33 catshark mandibles showing the dental lamina (yellow arrows) labeled with various A) phalloidin B) anti-pan actin. Scalebar: 100 µm.

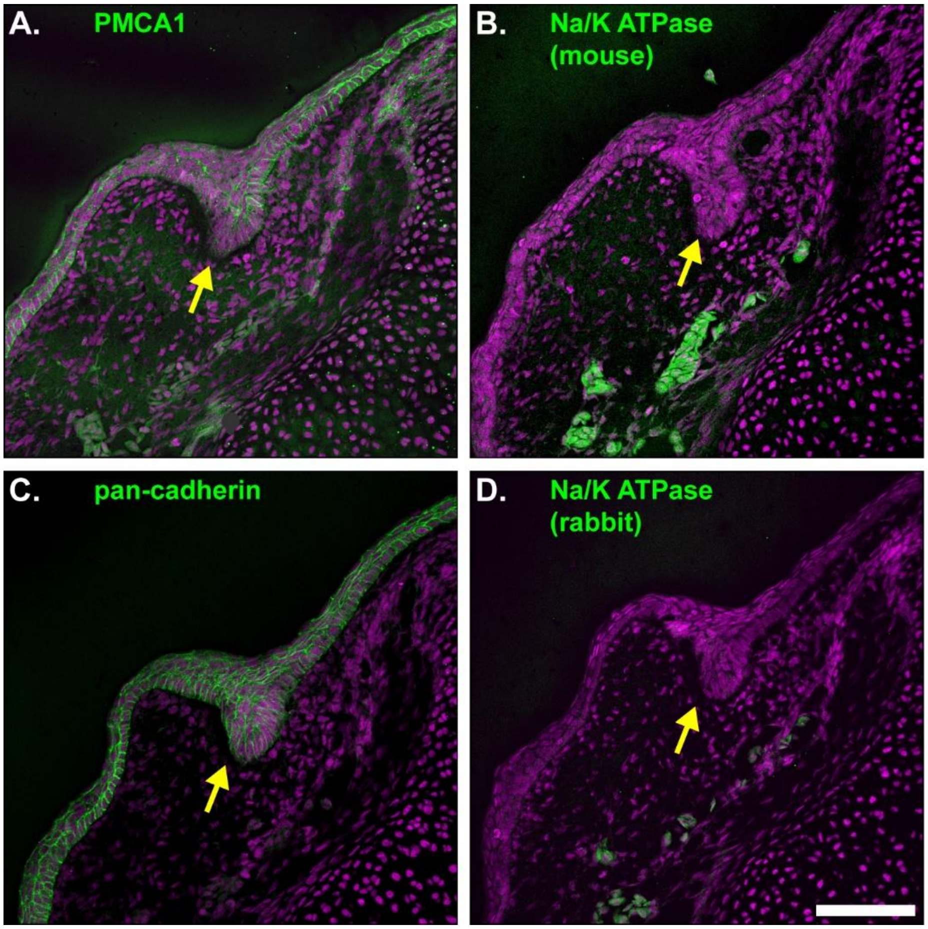


**Figure S3**. Representative serial thin sections of stage 32 catshark mandibles showing the dental lamina (yellow arrows) labeled with various plasma membrane antibodies, counterstained with DAPI (magenta). A) Monoclonal rabbit anti-PMCA1, B) monoclonal mouse anti-alpha 1 sodium potassium ATPase, C) monoclonal rabbit anti-pan cadherin antibody, intercellular junction Marker, and D) monoclonal rabbit anti-alpha 1sodium potassium ATPase. Scalebar: 100 µm.

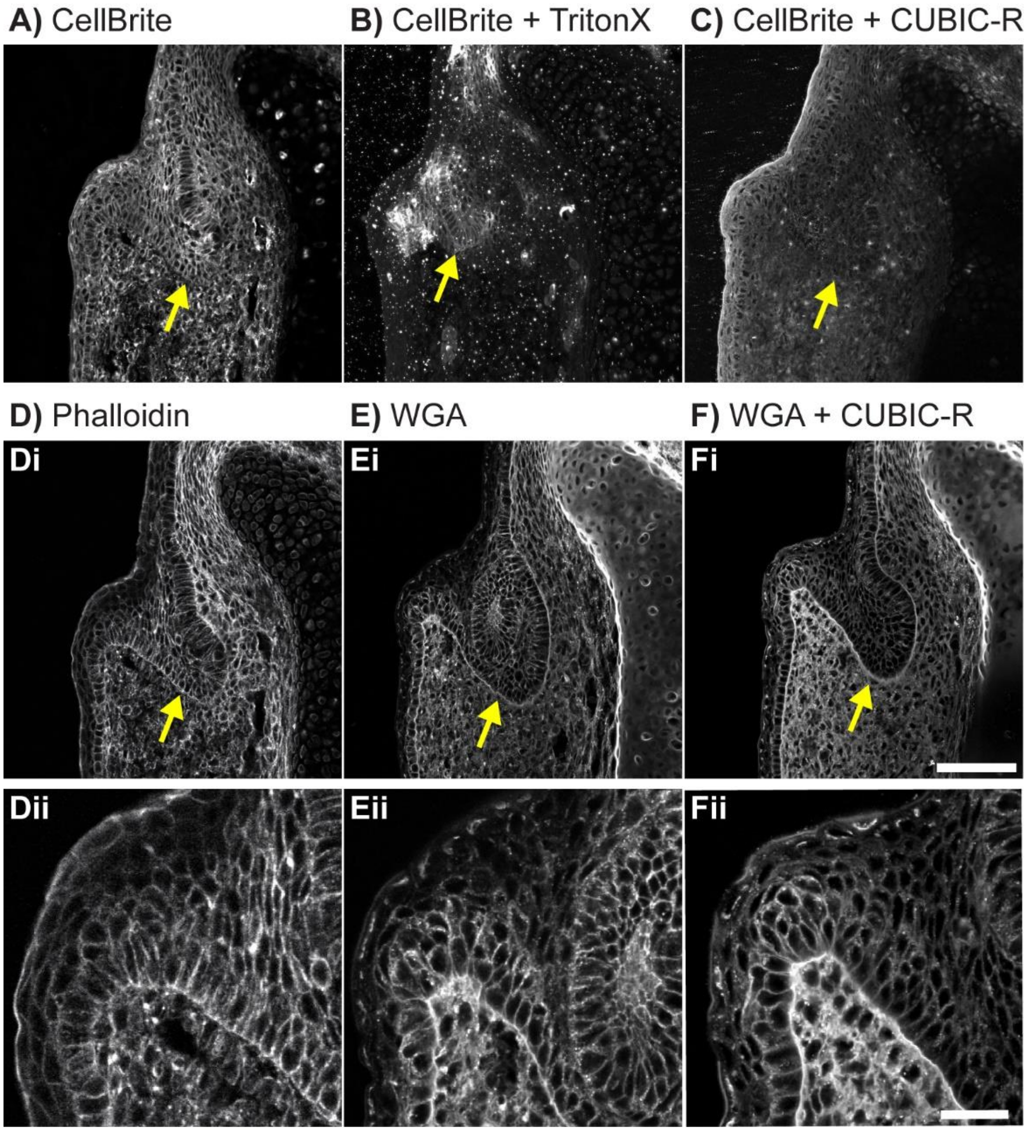


**Figure S4**. Representative serial thin sections of stage 33 catshark mandibles showing the dental lamina (yellow arrows) labeled with different cell boundary markers. Sections were labeled with CellBrite A) without Triton-X or CUBIC-R treatment; B) permeabilized with 0.1% Triton-X; or C) mounted with CUBIC-R. D) Section labeled with phalloidin and mounted with PBS. Sections labeled with wheat germ agglutinin (WGA) and mounted with E) PBS or F) CUBIC-R. Dii-Fii) Respective zoomed-in images. Scalebars: A-C, Di-Fi) 100 µm; Dii-Fii) 30 µm.

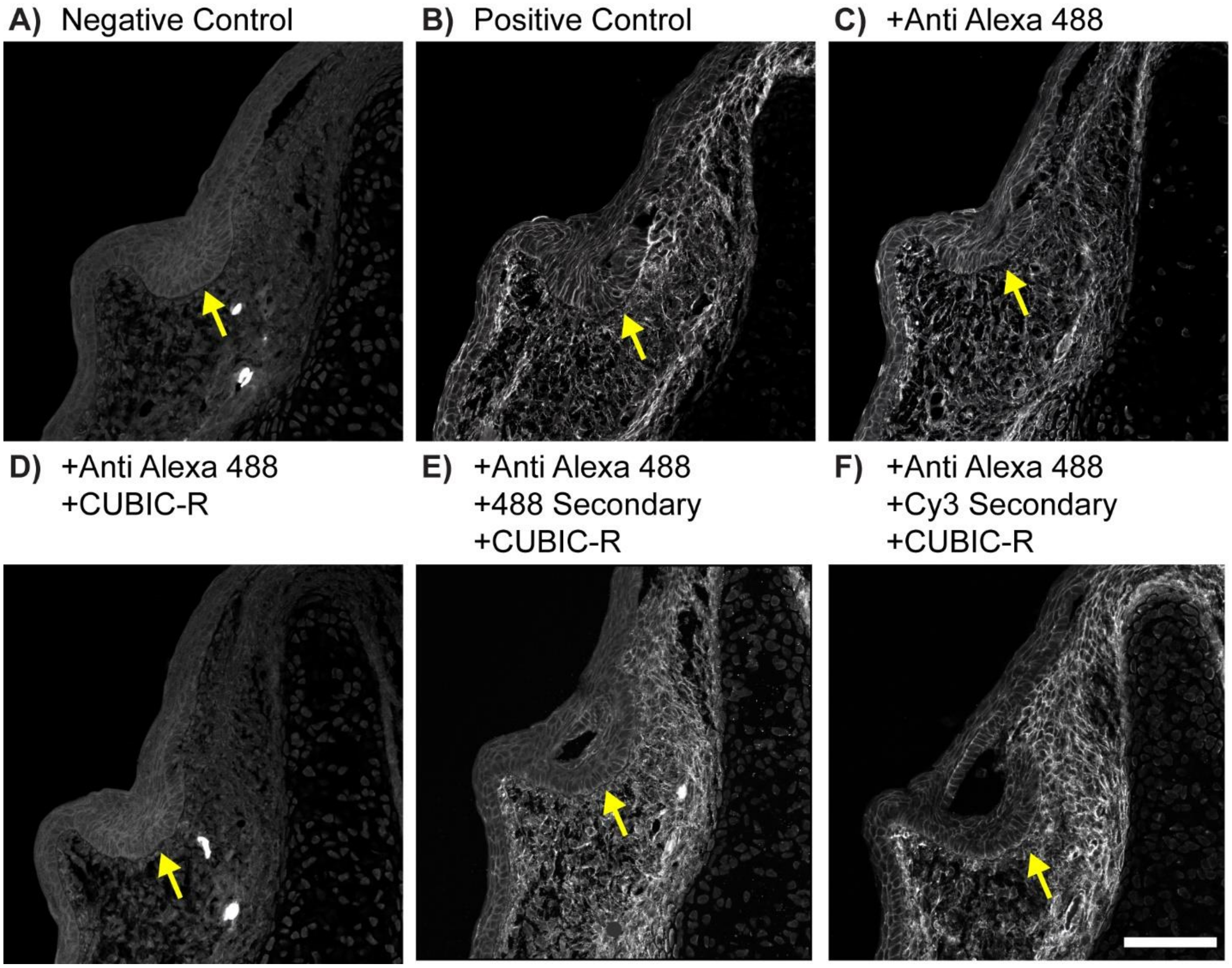


**Figure S5**. Representative serial thin sections of stage 32 catshark mandibles showing the dental lamina (yellow arrows) and labeled with phalloidin. Phalloidin-Alexa Fluor 488 labeled section mounted in A) CUBIC-R or B) PBS. Phalloidin-Alexa Fluor 488 labeled section incubated in anti-Alexa Fluor 488 primary antibodies and mounted in C) PBS and D) CUBIC-R. Phalloidin-Alexa Fluor 488 labeled section incubated in anti-Alexa Fluor 488 primary antibodies and E) Alexa Fluor 488 and C) Cy5 secondary antibodies then mounted in CUBIC-R. Scalebar: 100 µm.

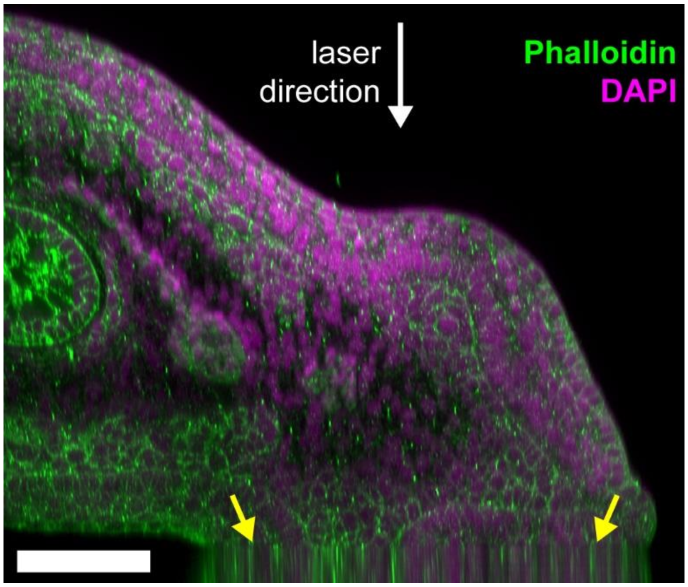


**Figure S6.** Example virtual section from 3D confocal volume of 32 mm catshark tissue with phalloidin retention protocol, counterstained with DAPI, and optically cleared with CUBIC-R refractive index matching media. Yellow arrows indicate region where objective has bottomed out and cannot image tissue at deeper penetration depths. Scalebar: 100 µm.

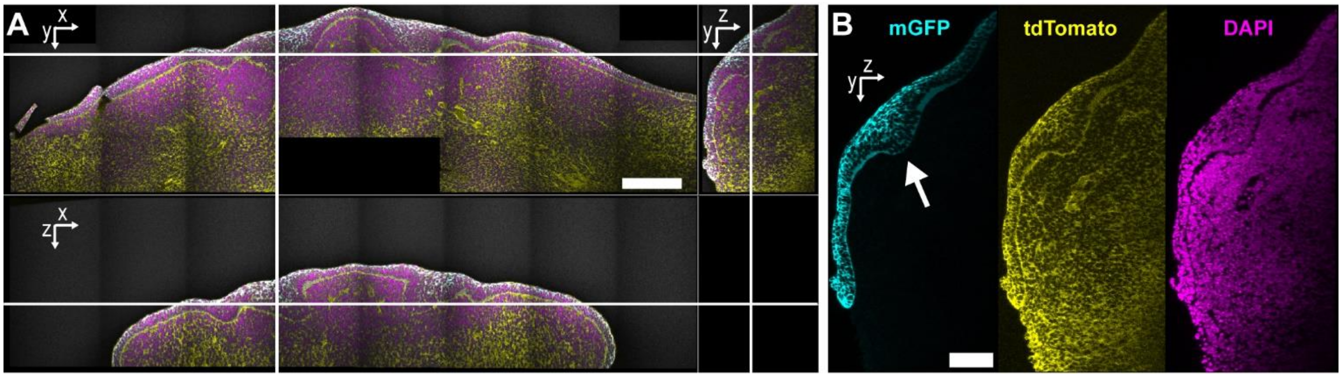


**Figure S7.** Example 3D confocal volume of E12.5 *K14*$^{Cre}$;*R26*$^{mT/mG}$ mouse mandible showing the dental lamina (white arrow) counterstained with DAPI and phalloidin-Alexa Fluor 633, polyacrylamide gel-embedded, and digested with proteinase K. A) Orthogonal views of 3D confocal volume. B) Single channel images of sagittal section. Scalebars: A) 200 µm, B) 100 µm.

**Video 1**. 3D Confocal volume of E12.5 *K14*$^{Cre}$;*R26*$^{mT/mG}$ mouse mandible cleared with CUBIC-R. Cells are labelled by membrane GFP (mGFP, cyan), mesenchymal cells retain expression of membrane tdTomato (yellow). Animation shows the orthogonal planes of the volume.

**Video 2**. 3D confocal volume of optically cleared 27 mm stage 29 catshark tail tip with phalloidin retention protocols applied, counterstained with DAPI. Animation scrolls through the sagittal slices of the 3D volume.

**Video 3**. Animated version of Figure 4. Example tail tissue from a 24 mm (stage 27) shark cleared and imaged to obtain individual 3D volumetric cell morphology properties. Volumetric images of A) input phalloidin fluorescence signal, and B) individually segmented cells. C-H) Heat maps of cellular morphology and phalloidin intensity. C) Average phalloidin intensity per cell, D) length of longest principal axis of each cell, E) cell sphericity, F) ratio of the largest/smallest principal axes, G) local cell density in units of cells/volume, and H) cell volume.
